# Working memory by distributed neural oscillators in a simple nervous system

**DOI:** 10.1101/2024.08.11.607402

**Authors:** Raymond L. Dunn, Caitriona M. Costello, Jackson M. Borchardt, Daniel Y. Sprague, Grace C. Chiu, Julia M. Miller, Noelle D. L’Etoile, Saul Kato

**Affiliations:** Weill Institute of Neurosciences, University of California San Francisco; San Francisco, CA, USA; Department of Cell and Tissue Biology, University of California San Francisco; San Francisco, CA, USA

## Abstract

Working memory allows an animal to gather sensory evidence over time, integrate it with evolving internal needs, and make informed decisions about when and how to act. Simple nervous systems enable careful mechanistic dissection of neuronal micro-dynamics underlying putative conserved mechanisms of cognitive function. In this study, we show that the nematode *C. elegans* makes sensory-guided turns while foraging and can maintain a working memory of sensory activation prior to the execution of a turn. This information is integrated with body posture to localize appetitive stimuli. Using a virtual-reality whole-brain imaging and neural perturbation system, we find that this working memory is implemented by the coupled oscillations of two distributed neural motor command complexes. One complex decouples from motor output after sensory evidence accumulation, exhibits persistent oscillatory dynamics, and initiates turn execution. The second complex serves as a reference timer. We propose that the implementation of working memory via internalization of motor oscillations could represent the evolutionary origin of internal neural processing, i.e. thought, and a foundation of higher cognition.

## Introduction

Any animal navigating a complex environment stands to benefit from the ability to quickly form impressions of the world, retain them internally, and act on them later. Such a working memory system allows for deferred, contingent actions, which may serve the animal better than reflex actions by allowing for more nuanced behavior or cognitive processing. But how did nervous systems acquire this ability? Distributed brain oscillations and their interactions have long been hypothesized to be a building block of cognitive function in complex-brained animals (*1*). Considerable theoretical work has been devoted to understanding how these observed phenomena might implement various cognitive functions (*2*). However, experimental establishment of causal roles and mechanistic understanding has been elusive, due at least in part to the sheer complexity of mammalian brains and the challenge of observing these neural networks at sufficient sampling density or completeness.

By contrast, in simpler brained animals with vastly lower neuron counts, oscillators serving to produce repetitive bodily movement, i.e. central pattern generators, have been closely studied and systematically dissected, yielding mechanistic insight in the production of adaptive but robust rhythmic motor patterns (*3*). But cognitive functions such as sensory-driven decision-making have rarely been studied in simple animals (*4*).

The 1 mm long nematode *C. elegans* crawls on its right or left side when on a flat surface. During foraging, the worm crawls in a straight or curved forward direction and punctuates bouts of forward crawling with discrete reversal-then-turn reorientation maneuvers, known as pirouettes, either in the dorsal or ventral direction (*5*, Fig. 1A). The choice of this direction has traditionally been described as an unregulated random process, akin to the randomizing tumbles of a bacterium performing “biased-random-walk” chemotaxis (*5*). However, worm olfactory neurons signal reliably on fast timescales (*6*), and mutations which slow temporal dynamics of olfactory receptors drastically disrupt chemotaxis, suggesting worms can integrate sensory information with instantaneous body posture.

**Fig. 1.**
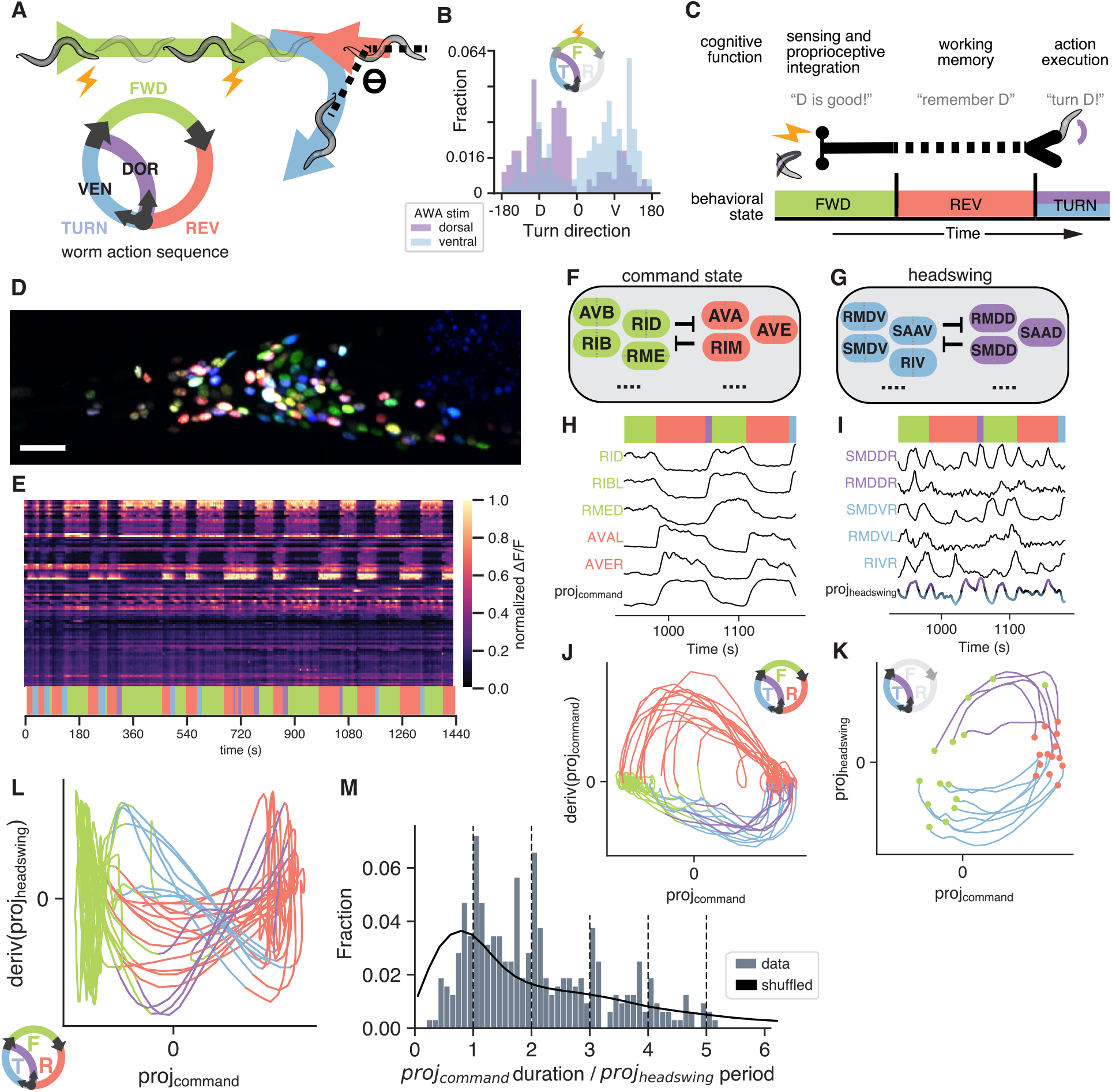
Worms make directed turns in a virtual olfactory environment, and discovery of a distributed headswing complex in whole-brain dynamics in paralyzed worms. **(A)** Schematic of closed-loop optogenetic stimulation of attractant sensory neuron with respect to instantaneous head curvature during forward crawling. [Inset] stereotypical motor command state sequence during foraging. **(B)** Angle of post-reversal turn following closed-loop stimulation, colored by whether stimulation was triggered by dorsal or ventral head bending during preceding bout of forward crawling. Test statistic (Tstat) = 77.32, p<0.0001. **(C)** Schema for deferred actions as a model for proto cognitive behavior. Memory of sensory experience is maintained across the reversal interval to turn execution. **(D)** NeuroPAL and pan-neuronal nuclear-localized GCaMP6s enable whole-brain imaging at cellular resolution with unambiguous neuron identification in restrained worms. Scale bar 20um. **(E)** Example heat plot of fluorescence (ΔF/F) time series of 109 segmented head neurons, one neuron per row. Top horizontal bars specify time intervals in (F, G). Bottom colored bar specifies the inferred command state. **(F-G)** Diagram of neuron classes with high magnitude in what we term the command state projection, which encodes FWD/REV, or the headswing projection, which encodes head curvature commands. **(H-L)** Representative neuron traces and phase plots from (E) of proj(command) or proj(headswing), and several neurons which participate in these complexes. Trajectories in (K) go from termination of preceding REV (red), until initiation of subsequent FWD (green) state. In trajectories of deriv(proj(headswing) vs. proj(command) state space (L), we see that there is a relationship between the onset timing, or phase of the post-reversal turn and the turn identity. **(M)** Distribution of proj(command) states (FWD or REV epochs), expressed in terms of contemporaneous headswing oscillation period. Shuffled (i.e. proj(command) and proj(headswing) periods are selected from random recordings) data do not show phase nesting. Tstat = 0.15, p<0.0001.

Here, we construct a closed-loop virtual olfactory environment (Fig. 1A,B), and show that worms integrate olfactory sensory dynamics with instantaneous body posture to navigate towards attractants, by executing directed turns that are deferred by tens of seconds, suggesting the existence of a working memory system. By monitoring neural activity across the brain, we find that this working memory is implemented by the phasic interaction of two distributed dynamical complexes associated with motor control. We contend that the internalization of motor command dynamics to direct deferred contingent actions may represent the evolutionary transition from reflex actions to actions guided by internal neural processing.

## Results

### C. elegans executes directed, sensory-guided reorientations

We engineered a closed-loop virtual odor environment to emulate a directional odor signal from the worm’s frame of reference by delivering optogenetic stimulation timed to particular body postures of the worm while crawling on an agar surface. We optogenetically stimulated the AWA sensory neuron, known to mediate chemotaxis to certain attractants, synchronized to either ventral or dorsal head swings during forward motion (Fig. 1A). In the absence of explicit sensory cues, when crawling on an agar surface, worm post-reversal turns are biased 60-75% towards their ventral side (*7*). After posture-timed optogenetic stimulation during bouts of forward crawling, individuals showed a strong preference for resolving the following reversal bout with a turn in the favored direction before resuming forward locomotion (Fig. 1B).

This observation suggests two key features of the sensory control of the behavior. First, to assign the stimulus to a particular spatial direction, the animal needs to integrate olfactory sensation with proprioceptive signals or efference signals of motor commands for head swings. Second, to turn in a favorable direction after an intervening reversal, the animal needs some form of working memory (Fig. 1C).

### Distributed dynamical complexes encode motor command state and head curvature

Studies across the animal kingdom have established a universal phenomenon that high-level motor commands are encoded in the low-dimensional state space of broadly distributed neural dynamics (4, 8-12). We asked whether lower-level sub-commands may also be encoded across many neurons, such as the sinusoidal movement of head swings in C. elegans. Previous studies have observed several neurons which correlate with dorsal or ventral head bending, however to what extent this signal reflects head musculature proprioception, motor output, causal decision processes, or any combination thereof, is unresolved (8, 13-16).

To look for neurons encoding headswing commands while avoiding the potential confound of proprioceptive state encoding, we restrained worms in microfluidic chips that enable high-quality volumetric whole-brain calcium imaging at cellular resolution (Fig 1D,E). In all trials, we observed cycles of the fictive command state sequence (forward-backward-turn) widely distributed across many neurons. For convenience, we define command states (and corresponding fictive neural network states): forward (FWD), reversal (REV), and dorsal or ventral (DOR, VEN) turn (TURN). As reported previously, the first principal component of whole brain recordings reliably encodes the command state sequence (8, Fig. 1F,H,J). We refer to the neural sub-network which exhibits these distributed dynamics as the command state complex, and the first principal component of these dynamics as the command state projection. Since the command state signal dominates the variance of brain-wide neuronal activity, we then performed PCA on the residuals of the command state projection, restricting time series data to periods for fictive forward locomotion only, which yielded a second strong, stereotyped network oscillation (Fig. 1G,I,K).

We found this faster oscillating component to be distributed across 14-20 neurons in the head of the animal that are variously implicated in motor control of the head muscles, sensorimotor integration, and proprioception (14, 17, Fig. 1G,I,K, S1A). The neuron class we observed with highest component loading was SMDD, and its contralateral neuron class SMDV had a large negative loading. Previous studies have shown SMDV reliably correlates with ventral head curvature during forward crawling and ventral post-reversal turns, and SMDD correlates with dorsal turns (8, 13, 15, Fig. S1B,C). Their anti-phasic coupling during FWD suggests their activity corresponds to fictive dorsal-ventral head swings consistent with sinusoidal crawling during the forward command state. We name this component the headswing projection. While TURN trajectories of different directions are mixed together in a single bundle together when plotted in command projection phase space (Fig 1J), they strongly split when plotted in joint command/headswing state space (Fig. 1K,L, Fig. S2A-F).

We asked if there was a relationship between the oscillation periods of the two projections. Interestingly, we found that distribution of the duration of FWD/REV epochs of the command state projection was enriched at integer multiples of the headswing projection periods (Fig. 1L, 1M). This coupling of internal dynamics is consistent with the nesting of headswings within forward crawling bouts previously reported (15). The persistence of an interaction across multiple fictive headswing periods suggested to us a potential substrate for maintaining a stable value, i.e. a memory, via the phase of the headswing oscillation, that could be leveraged for executing deferred sensory-guided decisions. To test this hypothesis, we examined neural network dynamics while evoking directed TURN.

### Decoding deferred, sensory-driven action in immobilized worms

We next combined closed-loop perturbative optogenetic stimulation with whole-brain imaging (Fig. 2A-C), recapitulating our virtual olfactory setup in immobilized worms. Each 16-24 minute recording captured 10-30 FWD-REV-TURN sequences. Within a trial, repeated AWA stimulations of ∼4-8s were delivered at a particular phase of the headswing projection oscillation (Fig. 2C,D).

**Fig. 2.**
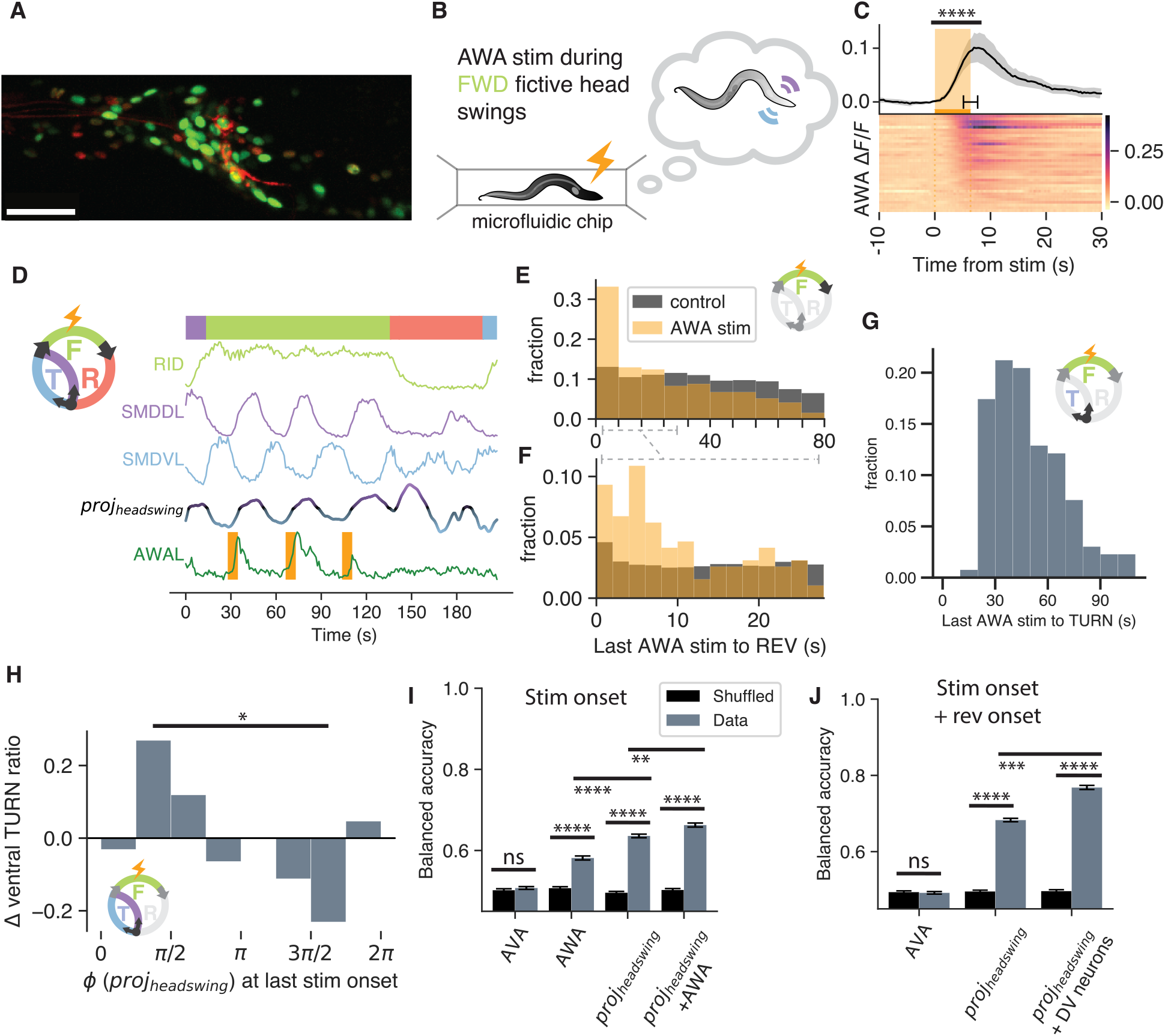
Decoding deferred, sensory-driven action in immobilized worms. **(A)** Maximum intensity projection of panneuronal GCaMP with WOrMsChRmine expressed under an AWA-specific promoter for acute, transient activation during whole-brain imaging. **(B)** Diagram of paralyzed worm under whole-brain volumetric calcium imaging with brain-state dependent AWA stimulation. Additional red expression is due to panneuronal tag-RFP present in NeuroPAL. **(C)** Heatmap of sorted individual AWA ∼4-8s stimulations during whole-brain imaging. Each row represents a single AWA from a different worm, and trial average is shown above. Tstat=-8.12, p<0.0001. **(D)** Example trial of simultaneous whole-brain imaging and closed-loop stimulation in paralyzed worms, recapitulating our simulated virtual olfactory environment in Fig. 1A. Olfactory activation was repeatedly embedded within spontaneous head-swing dynamics during forward crawling. Upon reversal initiation, stimulation was then withheld. **(E-F)** Distribution of time after AWA stimulation until next fictive reversal (REV), compared to spontaneous state transition computational control. Tstat=0.29, p<0.0001. **(G)** Distribution of time after AWA stimulation until dorsal or ventral TURN. **(H)** Change in ventral TURNs ratio, relative to unstimulated control and stratified by phase of proj(headswing) at the time of stimulation onset. A logistic regression model predicting turn direction based on the sine of phase was significant (LLR p=0.018, n=132). The fitted model showed that dorsal turns were more likely near 3π/2 and ventral turns were more likely near π/2 (β = -0.724, z = -2.294, p = 0.022). **(I)** Decoder balanced accuracy for TURN choice (VEN or DOR). Decoder was fit to neural activity at stimulation onset of AVA, AWA, proj(DOR/VEN), or AWA+DOR/VEN. Error bars are SEM after cross validation. For comparisons in (I,J), please see Table S1. **(J)** Same as in (I) but for decoder fit on features from both stimulation and reversal onsets.

Stimulation of AWA shifted the distribution of times until the next REV initiation (Fig. 2E,F), sometimes eliciting early reversals. We observed a secondary bump in the shifted distribution of time-to-REV for AWA stimulations, at ∼20s, which roughly corresponds to one typical period of headswing oscillation under immobilization (Fig. 1I, 2D), suggesting that we may observe the system as it maintains a sensory-evoked memory trace over the course of one of more fictive headswing oscillations.

We evaluated the direction of the TURN produced after the intervening REV state, about 30 to 90 seconds in the future (Fig. 2G), and found that our stimulations biased the DOR/VEN ratio in both directions, depending on the value of the estimated headswing phase at the time of last stimulation before a reversal (Fig. 2H).

Next, we trained a decoder on individual stimulation trials (see Methods) to predict TURN direction using activity leading up to the prior REV. We found that the dorsal/ventral identity of fictive post-reversal turns could be predicted by evoked AWA neural responses when combined with phase information of headswing projection at the time of stimulation (Fig. 2I), consistent with the observed effect of stimulation biasing turn choice (Fig. 2H). Prediction improved by including headswing state at REV onset (Fig. 2J), suggesting a post-stimulation role for headswing state in informing the deferred turn decision. Including features of individual headswing complex neuronal activity alongside the headswing projection further increased prediction performance, possibly suggesting that linear projection of headswing network dynamics may not fully capture the network dynamics underlying the sensation-to-action process.

The ability to predict deferred fictive turn decisions from prior neural activity in an immobilized, paralyzed setup demonstrates the existence of an internal neural working memory. Given the evidence of interaction between headswing projection oscillation and command state oscillation periods, we sought a closer study of oscillatory headswing dynamics during the intervening reversal.

### Internalized headswing oscillation phase is set after REV onset

During reverse locomotion, the head oscillations present during forward crawling are largely suppressed – the animal’s head trails behind the body (*18*). We found that during REV epochs in paralyzed worms, headswing projection dynamics continue to display oscillatory behavior; however, the magnitude is typically attenuated and the regularity of the oscillation waveform appears reduced (Fig. 1G, 2D, S2D,E). We refer to these dynamics as internalized because they persist in the absence of both sensory input (i.e. in completely immobilized worms) and motor behavioral output (i.e. during reversal/REV). Toward the end of a reversal epoch, the magnitude of the headswing oscillation typically increases over the course of 1-2 oscillations, exhibited in SMDD/V as well as other neurons. These data suggest that the headswing projection oscillation decouples from motor output during reversals but does not disappear, and recouples leading up to a reversal termination in order to effect the post-reversal turn.

To compare headswing oscillation amplitude, frequency, and phase during FWD and the following REV epoch, we fit single sinusoids to smoothed derivatives of the headswing projection as well as a set of high-loading neurons which participate in the headswing complex (Fig. 3A, Fig. S3). We observed statistically significant correlations between amplitude and frequency between headswing during FWD and REV (Fig 3B,C), suggesting they may be stable system properties independent of actual movement production, but we found no correlation between phase during FWD and REV, suggesting reset of phase (Fig 3D).

**Fig. 3.**
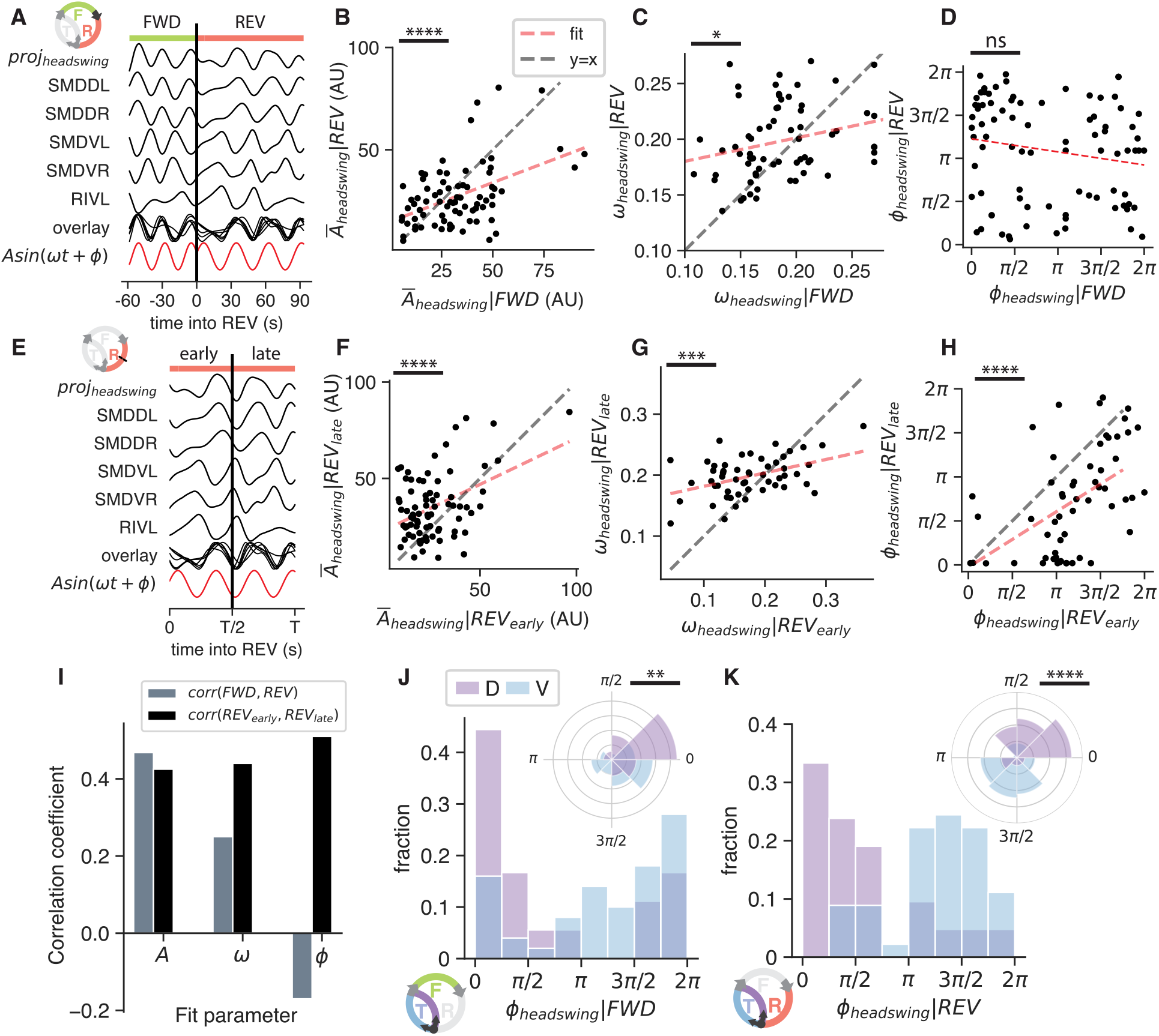
Internalized headswing oscillation phase is reset at REV onset. **(A)** Schematic of functional form fitting. A single sine wave was fit to the smoothed derivative of proj(headswing) as well as high-loading individual neurons composing it, for each paired FWD and subsequent REV segment. By convention, the sign of dorsal-associated neuron loadings was fixed to be positive. Sines were fit on continuous time coordinates for direct comparison of fit parameters. **(B-D)** Correlation between sine wave neuron loading (B), frequency (C) and phase (D) in FWD and corresponding REV segment. **(E)** Schematic for fitting sine separately to first (early) and second (late) halves of a REV segment. **(F-H)** Correlation between neuron loading onto the sine, frequency, and phase between first and second halves of a REV segment. **(I)** Summary of correlation coefficients calculated in (B-D, F-H), see Table S1 for each comparison. (J-L) Sine wave phase in FWD and REV segments, colored by future TURN direction.

Despite the irregularity and reduced magnitude of the headswing projection during the initial segment of a REV epoch, we wondered if the headswing complex maintained the ability to convey information in its oscillatory activity until the time of REV termination. We again fit sinusoids to activity during REV; however, this time we separately fit on the first and second halves of REV (Fig. 3E) and compared fit parameters. We found small but statistically significant correlations in amplitude, frequency, and phase between early and late REV, suggesting that the internalized headswing oscillation can be stable over the course of a REV epoch (Fig. 3F-H).

Returning to headswing dynamics during FWD or REV segments, we found that the phase of our sinusoid fits stratified turn choice, especially during REV (Fig. 3J,K). Therefore, we conclude that the phase of the headswing complex is clustered to one of two intervals [0, π] or [π, 2π] when internalized to store the intent of future turns.

### Headswing-complex neurons terminate the REV command state

While the head projection oscillation is distributed across many neurons and is implicated in the memory process, it is possible that the locus of the memory that is causal for action is limited to a subset of neurons or even one neuron. To probe the sufficiency of individual neurons in holding a sensory memory or driving turn identity, we selectively stimulated headswing complex neurons. We used a digital micromirror device to spatially restrict our laser stimulation to selectively stimulate either SMDD or SMDV neurons, timing stimulations to occur during fictive reversals (Fig. 4A,B). Surprisingly, selective optogenetic stimulation, though demonstrably depolarizing the neurons (SMDV shown in Fig. 4B,L), did not affect the resulting TURN direction (Fig. 4M, Fig. S4A). This suggests that the activity of SMD neurons, while clearly signaling TURN identity, are not the causal locus of the memory trace. Interestingly, despite the inability to bias turn direction, stimulation of either SMD neuron class often elicited immediate reversal terminations (Fig. 4C,D,H).

**Fig. 4.**
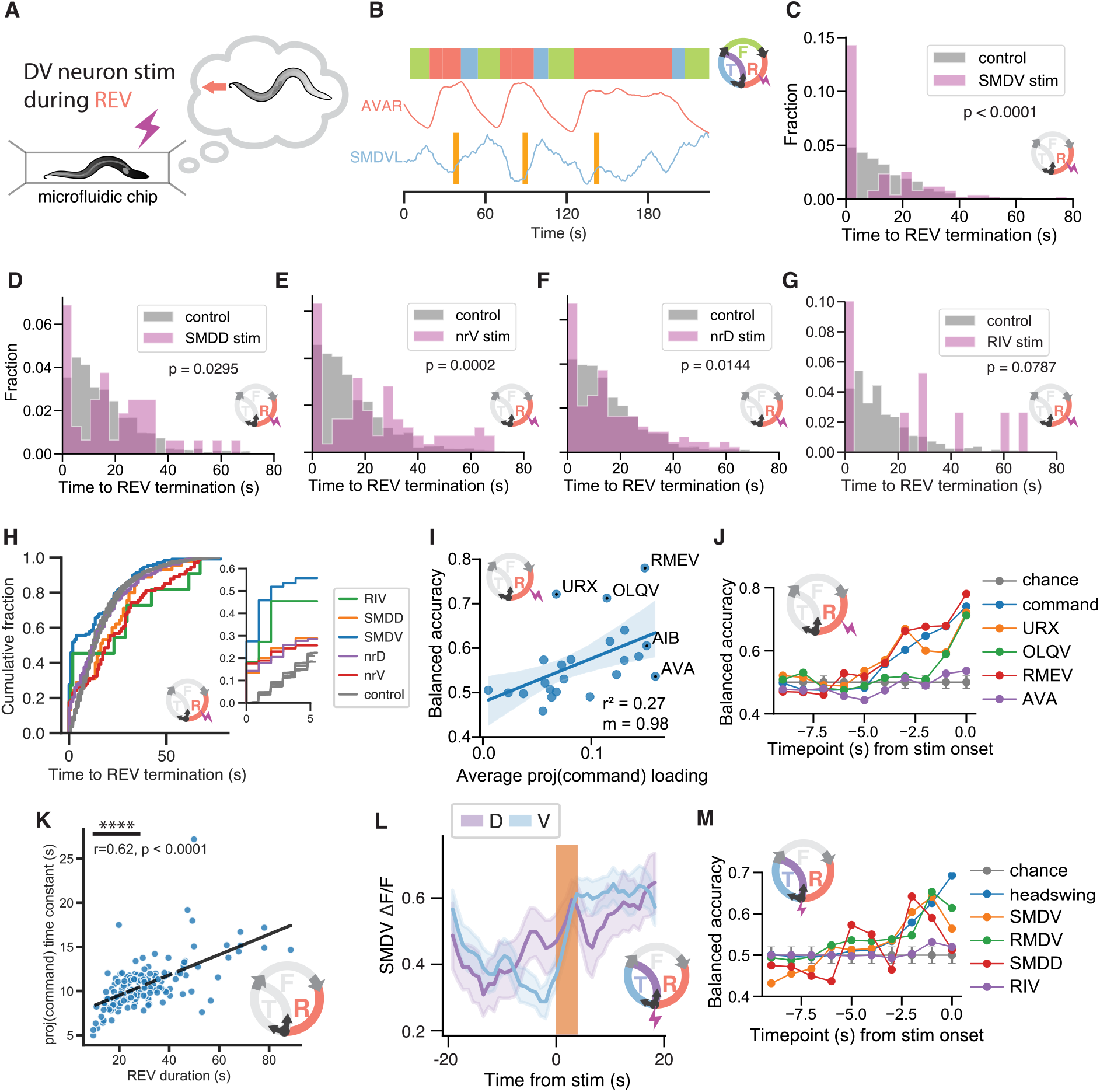
Headswing-complex neurons terminate the REV command state. **(A)** Diagram of paralyzed worm under whole-brain volumetric calcium imaging with brain-state dependent activation of neurons with high magnitude loading in the headswing complex. **(B)** Example trial of single neuron optogenetics targeting SMDV timed to fictive reversals, denoted by AVA depolarization. **(C-G)** Distribution of time after stimulation until REV termination following stimulation of headswing neurons (SMDV/SMDD/RIV or nrV/nrD subcompartments of RIA). Computational control for spontaneous state transition is shown in grey. (H) Empirical cumulative distributions detailing the effect of stimulating various DOR/VEN neurons (SMDV/SMDD/RIV or nrV/nrD subcompartments of RIA) on time after stimulation until REV termination. Inset: expansion of timescale immediately after stimulation. See Table S1 for each statistical comparison. **(I)** Balanced accuracy of decoding REV termination after activation of RIA nrV or nrD, based on neuron features at the time of stimulation correlates with proj(command state) loading. **(J)** Balanced accuracy of decoding successful REV termination, by fitting a decoder on activity prior to stimulation. Same data as (I). **(K)** The rise constant of the REV command state correlates with the reversal duration (r=0.62). **(L)** Average evoked SMDV activity on trials where stimulation was immediately followed by dorsal (purple) or ventral (blue) TURN. Stimulation had no effect on future turn identity (see Supplementary Information). Shaded regions indicated 95% CI. Note that given future D/V turn identity, SMDV activity has already diverged prior to stimulation. Following stimulation, SMDV shows prolonged and sustained depolarization for ventral but not dorsal TURNs, where depolarization is confined to light-on period. **(M)** Balanced accuracy of decoding TURN direction following activation of RIA nrV or nrD, by fitting a classifier on activity of various neurons features prior to stimulation.

We next turned to the head neurons RIA and RIV. Previous studies have implicated RIA in sensory-proprioceptive integration; it exhibits compartmentalized calcium dynamics along the nrV and nrD regions of its nerve ring neurite, which tightly correlate with SMDV and SMDD activation during ventral and dorsal head bending, respectively (*13*). The motor neuron RIV has high correlation with SMDV but few shared synaptic outputs (*17, 19*). We found that spatially localized optogenetic stimulations of sub-compartments of RIA, and as well as RIV, also immediately terminated reversals (Fig. 4E-G). Previous studies have observed that chronic inhibition of SMDs or optogenetic activation of SAAV and SAAD, which correlate with head curvature during reverse crawling, can impact reversal timing as well (*15, 20*). The observations that 5 different tested neuron classes of the headswing complex can terminate reversals but those investigated do not bias turn direction suggest that the command intent of turn direction is maintained by a dynamical complex distributed across several neurons.

### Command state dynamics serve as a reference clock for headswing oscillations

How does the animal know when to recouple the internalized headswing oscillation and terminate a reversal to produce the correct turn direction? To store a persistent value (i.e. memory) reliably using the phase of an oscillator over several oscillation periods, a reference clock with a predictable relationship to the oscillator is needed. Interestingly, the capability of RIA subcompartments to terminate fictive reversal was gated by the instantaneous activity of the command state projection at the time of stimulation on a per-trial basis (Fig. 4I,J). Furthermore, individual neuron predictivity was positively correlated with command state projection loading (Fig. 4I,J). One interpretation of the ability of headswing neurons to immediately terminate fictive reversals is proprioceptive; they signal the successful execution of physical bending that produces reverse movement. As the command projection evolves, the system becomes permissive for proprioception-driven reversal termination. We fit an exponential to the command projection during REV (Fig. 4K, Fig. S4B) and found that the time constant of command projection ramping is correlated with reversal duration, the timescale over which the memory must be maintained, suggesting that the command complex may function as a reliable time reference. The decay in activity of positive loading neurons such as OLQ, URX, AIB or the ramping of neurons with negative loading such as RME, could potentially function as behavioral timers in conjunction with other sensory or motor roles previously proposed (*21-23*).

Focusing next on stimulations that elicited a TURN by terminating REV, we found that the activity level of the headswing projection, or alternatively individual neurons with high headswing loadings, was predictive of the TURN direction (Fig. 4L,M). Using our decoding framework, we were unable to find neurons whose static, monotonic, or differential activity levels were predictive of turn identity. We suspect the non-monotonic time course of predictivity is a consequence of oscillatory dynamics in these neurons; the headswing projection captures both DOR and VEN depolarizations, whereas individual headswing complex neurons display strongly turn-direction-dependent differential activity only during half of their oscillation cycle. The simultaneous ramping of the command state complex and period nesting with the headswing complex (Fig. 1M) may allow the animal to balance two normative functions: drive to terminate reversal and turn in the correct direction.

### A dual half-center oscillator model reproduces nested projection dynamics

The emergence of stable, spontaneous oscillations from mutually inhibiting neurons or groups of neurons has been extensively modeled, exemplified by the half-center oscillator (HCO) (*24*, Fig. 5A). The HCO model produces push-pull oscillatory dynamics strongly reminiscent of the waveforms of the command state projections and headswing projections (Fig. 1H,I). We modeled command state and headswing projections as separate HCOs (Fig. 5B, see Methods), and found that the incorporation of an interaction term between the two HCOs, modeling the influence on command state projection by the headswing complex (Fig. 4), produced a nested distribution of state durations resembling the experimental distribution (Fig. 1M) when random current is injected into a unit, representing fluctuations from sensory input and other internal processes (Fig. 5C). This model raises an intriguing mechanistic hypothesis: the worm brain may be composed of multiple loosely coupled dynamical complexes (e.g. HCOs) whose coordination implements cognitive functions such as working memory and supports purposeful behavior.

**Fig. 5.**
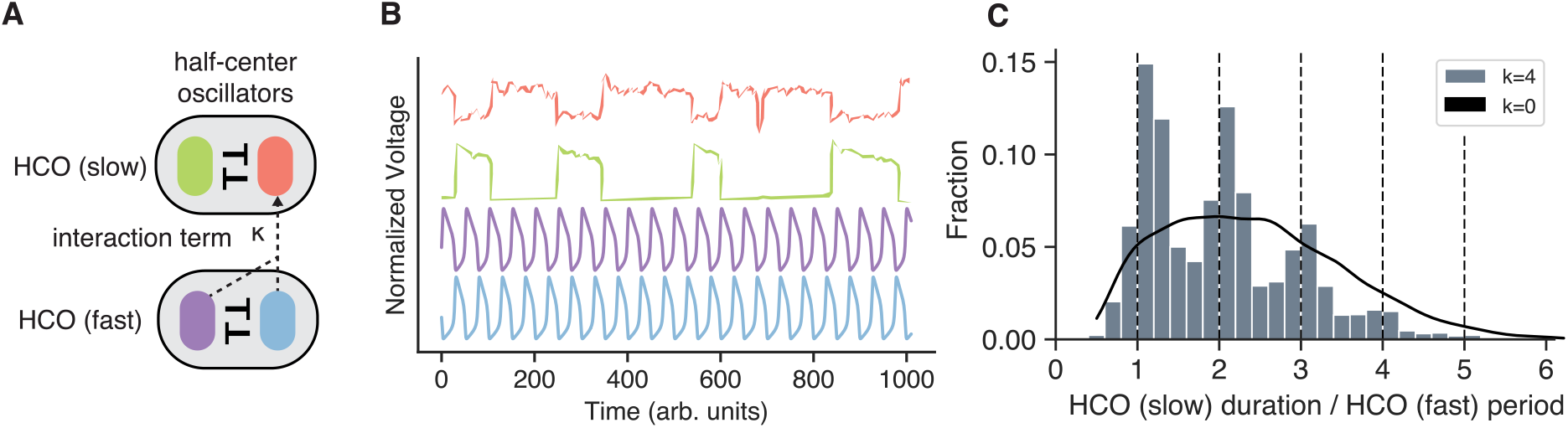
Dynamical model of interacting nested oscillators. **(A)** Diagram illustrating model components; two half-center oscillators (HCOs), and a coupling coefficient k reflecting the input from one HCO onto a neuron in the other. **(B)** Example parameterized dynamics of the Morris-Lecar neurons which compose each HCO. **(C)** Nonzero interaction between the two HCOs recapitulates state duration nesting reminiscent of Fig. 1M.

## Discussion

We found that worms are capable of producing directed turns by gathering sensory evidence and integrating it with body posture, revising a longstanding notion that foraging reorientations are undirected. Our data suggest that worms can navigate odor environments based on the following mechanism: (a) sensory input is received during crawling sinusoidal head swings and integrated with body posture to form sensory memory, (b) headswing and command state dynamics are coordinated, and headswing phase is set to the interval [0, π] or [π, 2π], (c) during reversal, a ramping process in the command state complex gates reversal termination, until (d) a proprioceptive signal indicates successful movement execution, terminating REV, and finally (e) a directed turn is produced based on whether the headswing complex is in DOR- or VEN-associated phase at the time of headswing recoupling.

Our experiments argue against the encoding of working memory in static neural activity levels for the sensory-guided decision-making capability. They instead suggest that working memory is implemented by the maintenance of the relative phase of two coordinated dynamical complexes.

We surmise that the original function of the head neural oscillator was to produce physical movement, and the motor-decoupled mode of operation only arose later to support the function of working memory, endowing the animal the ability to perform deferred sensory-guided action selection. This evolutionary step, from embodied to internal neural oscillations, may represent the origin of a functional primitive of cognition subsequently recruited to do far more complex forms of thinking.

The dynamical model we present in Figure 5 provides an expressive basis for building more sophisticated models. Features such as additional neural subnetworks or neuropeptide signaling could be incorporated to capture the cognitive processes enabling the worm’s full behavioral repertoire.

## Methods

### Behavior imaging and quantification

Worms were prepared 24 hours in advance of imaging: L4 worms were picked onto fresh NGM agar plates with an OP50 lawn. Immediately before imaging, worms were picked onto a fresh NGM agar plate without food, into a drop of M9. Single worms were aspirated out of that drop and washed 3x by aspirating into new drops of M9. Finally, the worm was aspirated onto an NGM agar plate without food, with care taken to re-aspirate excess M9. Worms were imaged while crawling on the surface of the agar without lid for 1-2 recordings of 12 minutes each. Data were acquired on an M205FCA motorized stereo microscope equipped with a LMT260 XY-Scanning stage, a TL5000 transmitted light base, an LED3 epifluorescence light source (Leica Microsystems) and combined with an ORCA-FLASH V3 sCMOS camera (Hamamatsu). Hardware was set up for control in micro-manager (25). Zoom was set to result in a field-of-view of 4.78mm. Worms were tracked over the course of an experiment with the microscope’s motorized stage centered on the worm’s centroid.

For experiments involving optogenetics, care was taken to reduce activation of the optogenetic channel from brightfield illumination light. About 24 hours prior to imaging, 80ul of 100uM all-trans retinal in M9 was added to the bacteria lawn of L4s. Worms were then placed in foil at 20C until imaging. During an imaging session, worms were partially covered if not being immediately prepared for an experiment. Optogenetic illumination was made in widefield mode with the LED3 set to 100%.

### Whole-brain Ca2+ imaging of C. elegans

Two-layer PDMS microfluidic devices were fabricated as described previously (*23, 26*). Worms were prepared 24 hours in advance of imaging: L4 worms were picked on to fresh NGM agar plates with an OP50 lawn. Immediately prior to imaging, worms were picked onto fresh NGM agar plates without food and placed in a drop of M9 solution with 5mM tetramisole. Worms were left in tetramisole for 10 minutes prior to being aspirated into the microfluidic channel. Worms were imaged for 16-24 minutes. Data were acquired on an inverted spinning disk microscope (Yokogawa W1-SoRa) set up on a DMI8 inverted microscope stand (Leica Microsystems) equipped with a Kinetix sCMOS camera (Telydyne Photometrics) and a Versalase Laser combiner (Vortran). Roughly half of recordings were acquired at 2×2 camera binning, the rest without binning. The microscope objective lens was 40x 1.25NA WI. Sample volumetric scans were performed using a piezo stage (ASI) with 10-12 z-planes with z-spacing 2.5-3um. In select recordings, 4 z-planes with z-spacing 3um were used to measure neurons in lateralized anterior, lateral, dorsal, and ventral ganglia at higher temporal resolution under equivalent optical conditions. Prior to acquiring videos for calcium timeseries, a reference high-resolution structural image was acquired using 40×1um z-planes under 4 different optical conditions to measure NeuroPAL fluorescence (*27*). Exposures ranged from 30ms - 80ms under 100 - 200uW/mm^2^ illumination to minimize photobleaching and optogenetics/fluorescence crosstalk.

For experiments involving optogenetics, care was taken to reduce activation of the optogenetic channel via illumination for measuring GCaMP fluorescence. About 24 hours prior to imaging, 80ul of 1uM ATR in M9 was added to the bacterial lawn of L4s. Worms were then placed in foil at 20 C until imaging. During an imaging session, worms were partially covered if not being immediately prepared for an experiment. Worms were imaged at roughly 70-140uW/mm2 (measured at sample plane) of light at the sample, for 20-30ms exposures within an 80 ms duty cycle - 20 ms-30 ms of light exposure and 50-60 ms of blank time. Optogenetic illumination was perfomed using a digital micromirror device (Mightex Polygon P1000) combined with an LDI-7 laser combiner (89North)) at 640 nm, powered to 10-20% of maximum laser intensity. Illumination was performed in widefield for worms with optogenetic construct expressed under a single-neuron promoter, and localized to about 5×5 um for selective single-neuron illumination of optogenetic constructs driven by multi-neuronal promoters.

Behavioral decoding of whole-brain recordings was performed as previously described (*8*).

### Region of interest (ROI) detection in volumetric Ca2+ imaging data

ROI detection from neural timeseries videos was adapted from (*8*), implemented by the Napari (*28*) **eats-worm** plugin. Briefly, interframe motion was first registered using manual tracking (*29*). A reference ROI movie was then generated composed of each image plane by averaging successive blocks of 20-200 movie frames to reduce noise. Each frame of the reference was adaptively thresholded based on median image brightness, median filtered, then convolved with a gaussian kernel. Local maxima were found and merged if peaks were adjacent within a greedy threshold. For each ROI center, a surrounding region with radius 5-7 was defined, with overlapping adjacent regions excluded via Voronoi tessellation with area shrinkage of 0.5 pixels. ROIs in adject timepoints were linked via local greedy matching. Cells below detection threshold were extrapolated based on the motion of neighboring ROIs. Finally, time-varying multi-plane ROIs were adjoined based on overlap. Each neuron was manually inspected for artifacts and overlapping fluorescence with adjacent neurons by R.L.D. For high temporal resolution half-brain imaging experiments, images were compressed along the Z axis via maximum intensity projection, becoming 3D ROIs. Neural time series extraction was adapted from (*8*). Briefly, for each 4D ROI, a single-cell fluorescence intensity was computed taking the average of the brightest 30-60 voxels at every time point after subtracting z-plane specific background intensity. Background values were computed by averaging pixels not belonging to any ROI within a radius of 21 pixels. DF/F0 was computed for each neuron, with F0 as mean background fluorescence.

### Identification of neuron genetic identities

In each recording, we detected 45-140 neurons. Neurons were identified by assessing their anatomical position, relationship to surrounding neurons, and their established activity patterns. Furthermore the strains used for experiments here express the NeuroPAL genetic cassette (*27*), which uses a genetically defined combination of 4 fluorophores to discriminate neurons based on multi-color reporter expression. Neuron labels were assigned by hand, using NeuroPAL documentation for guidance. In many cases ambiguity still existed, so here we opted for a more conservative approach and chose not to ID neurons which could not be identified beyond reasonable doubt. In rare cases, ambiguous identities are denoted in parentheses.

### Neural time series derivatives and embeddings

Derivatives and PCA on neural time series data was performed as previously described (*8, 30*). Specifically, total-variation regularization was used to compute de-noised time derivatives while resolving the accuracy of command state transitions to single frames. Differentiation ensures some degree of stationarity to the signal, improving subsequent analyses. This approach was also used for the calculation of proj(headswing). In more detail, for calculating proj(headswing), we first subtracted the sums of projections to the derivative time series onto temporal PC1. Next, we sub-segment forward command states and remove unsustained forward states defined by a threshold (<4s). The resulting segments are concatenated for PCA (*31*) to compute loadings. Full time series are then projected onto these loadings. Prior to PCA, time series were detrended and regularized. For comparison across animals, we applied PC matching by inverting projections such that the genetically identified neuron SMDV would be negative. In some cases, such as a large number of neurons drifting out of the focal plane contributing high variance, this procedure was adjusted by first filtering neurons with thresholded (0.5) normalized covariance to SMDV or by cropping the beginning, or end, of the full time series.

### Linear predictive modeling framework

For predictive modeling, we used KNN and SVM classifiers, respectively, implemented by **scikit-learn** (*32*) with 20-fold cross-validation to limit overfitting. To account for class balance, we report the balanced accuracy metric, which is equivalent to accuracy score with class-balanced sample weights. Model performance was averaged over 100 iterations with randomized seeds for cross-validation. For timepoint predictive analysis, separate models were fit on neural activity (DF/F and derivative as independent features) at single timepoints relative to stimulation. In both cases, control distributions were generated by shuffling class labels.

### Evaluating the effect of neuron stimulation on global state transition

The significance of the effect of neuron stimulation on command state transitions (time to REV initiation or REV termination) was assessed using a Kolmogorov-Smirnov test. The comparable control distribution must account for spontaneous command state transitions unaffected by stimulation. To calculate control null distribution, first we took unstimulated states from control worms and computed the marginal distribution of time until state transition. We then accumulated shifted state durations selected from our experimental group, based on time from stimulation, to generate the marginal distribution of time until state transition.

### proj(command) ramping and phase analysis

The time constant of proj(command) ramping was calculated by fitting a saturated curve to proj(command) traces scaled to each reversal. proj(command) and proj(headswing) phase were calculated by smoothing both traces with a gaussian filter, identifying extrema (scipy.signal.find_peaks), and linearly interpolating between peaks and valleys. The ratio of proj(command) to proj(headswing) period was calculated using the inter-peak-intervals of each signal for the corresponding cycles.

### Function fitting

For fitting sinusoids to headswing-associated neurons, we selected a subset of neurons with high average embedding in proj(headswing) to fit on: SMDVL, SMDVR, SMDDL, SMDDR, RIVL, RIVR, and additionally proj(headswing) and fit the following functional form: *A sin*(*ωt*+*φ*), Where A is a vector of neuron weights, omega is the frequency of the sine, and phi is the phase offset. Parameters were fit with the Nelder optimization procedure using the python package lmfit. Minimum and maximum constraints were placed on fit parameters depending on the parameter being evaluated; for example, when evaluating omega or phi, dorsal and ventral head curvature associated neurons were constrained to have non-zero amplitudes of opposite sign and curves were normalized to the interval {-1, 1}. We further conducted these analyses on trials with and without AWA stimulation. For specifics, reference the accompanying code. In all cases, we fit on the numerical derivative of neuron curves, estimated with total-variance regularization (30), and these curves were then smoothened with a box filter. We found our results were similar without temporal smoothing. For parameter initial values, omega = 0.18, phi = pi, and amplitudes were as follows: proj(headswing), SMDDL, and SMDDR: 1, SMDVL, SMDVR: -1, RIVL, RIVR: -0.5.

### Dynamical model of nested oscillators

We implemented a model with two connected half-center oscillators (HCOs) described in (24). Each HCO is composed of two Morris-Lecar neurons (*33*), i.e. models of graded conductances. Each Morris-Lecar neuron consists of two voltage dependent conductances, a leak current, and an inhibitory synaptic current:

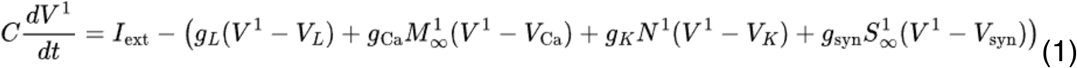

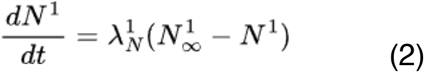

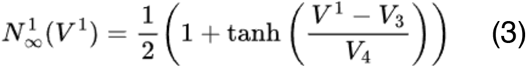

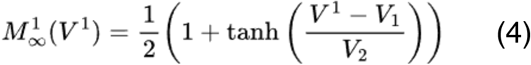

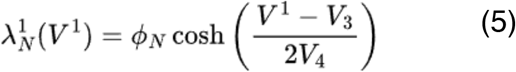

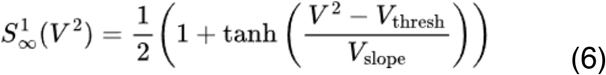

We parameterize two HCOs, one to model the dynamics seen in the command projection, and one for the headswing projection. We simulated this system using a 5th-order Runge-Kutta solver or, for expediency, the Euler method, which did not affect the qualitative results. In this study, we extend this model by adding a noise process and interaction term to the external applied current to one unit in the slow HCO:

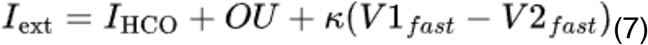

Where I(HCO) is external applied voltage, OU is an Ornstein–Uhlenbeck process, and V1 and V2 are the voltages of the two units in the fast HCO. Together these equations produce a distribution of spontaneous state durations, which exhibit nesting at k>0. For all model parameters, please see accompanying code.

### Table of statistical comparisons

Please see **Table S2** (Supplementary Information) for details of statistical comparisons made in this study.

### Data and materials availability

All data and code are available at WormID.org.

## Notes

## Acknowledgments

We thank members of the Kato and L’Etoile labs for useful discussion. Data for this study was acquired at the UCSF Innovation Core at the Weill Institute for Neurosciences on a custom imaging system controlled by the open-source software Micromanager. Special thanks to Dr. Caroline Mrejen for assistance with microscopy design and implementation. Thanks to Dr. Cori Bargmann for sharing the Chrimson plasmid. Thanks to Drs. Oliver Hobert and Evitar Yemini for sharing NeuroPAL strains.

## Author contributions

Conceptualization: RLD, SK

Resources: JMM, GCC

Formal Analysis: RLD, CMC, SK

Methodology: RLD, JB, SK

Investigation: RLD, DYS, GCC

Funding acquisition: RLD, NDL, SK

Writing – original draft: RLD, SK

Writing – review & editing: RLD, SK, CMC, NDL

## Funding

Some worm strains were provided by the CGC, which is funded by NIH Office of Research Infrastructure Programs (P40 OD010440).

Research funding was provided by National Institutes of Health, specifically R35GM124735

National Institutes of Health (SK) R35GM124735

National Institutes of Health (NDL) R01DC005991

National Institutes of Health (NDL) R01NS087544

National Institutes of Health (RLD) F31NS115572

Weill Institute for Neurosciences (SK)

Weill Neurohub (SK)

Chan Zuckerberg Initiative (JB)

## Supplementary Information

Supplemental information is available for this paper. See supplemental information for Figures S1-4, and Table S1-2.

