## Supplementary Information for "Working memory by distributed neural oscillators in a simple nervous system"

**Contents:**
Figs. S1 to S4
Tables S1 to S2

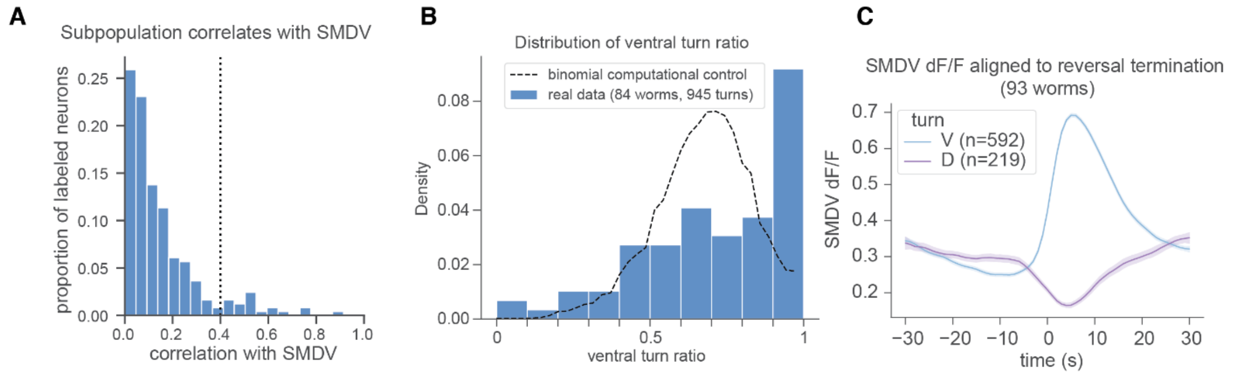


**Fig. S1. Ventral turn ratio, dynamics, and neural correlations with SMDV. (A)** Proportion of neurons labeled with NeuroPAL which exhibit high (>0.4) Pearson's correlation with SMDV. Correlation was calculated on normalized derivatives. The distribution of activity correlations with SMDV appear bimodal, which we interpret as a hint that headswing subcomplex dynamics may be a distinct dynamical entity. **(B)** Distribution of ventral turn ratio on a per-worm basis, computed from all post-REV TURNs. Deviation from the expected distribution of turn ratios if all turns were independent, computational control, which we interpret as a hint that turning selection may be under regulation. **(C)** Average activity of SMDV prior to REV termination. Shaded regions indicate 95% confidence intervals. Worms produced more fictive ventral turns on average. Prior to reversal initiation, SMDV is more hyperpolarized compared to when approaching fictive dorsal turns, which we interpret as a hint to intrinsic phasic dynamics.

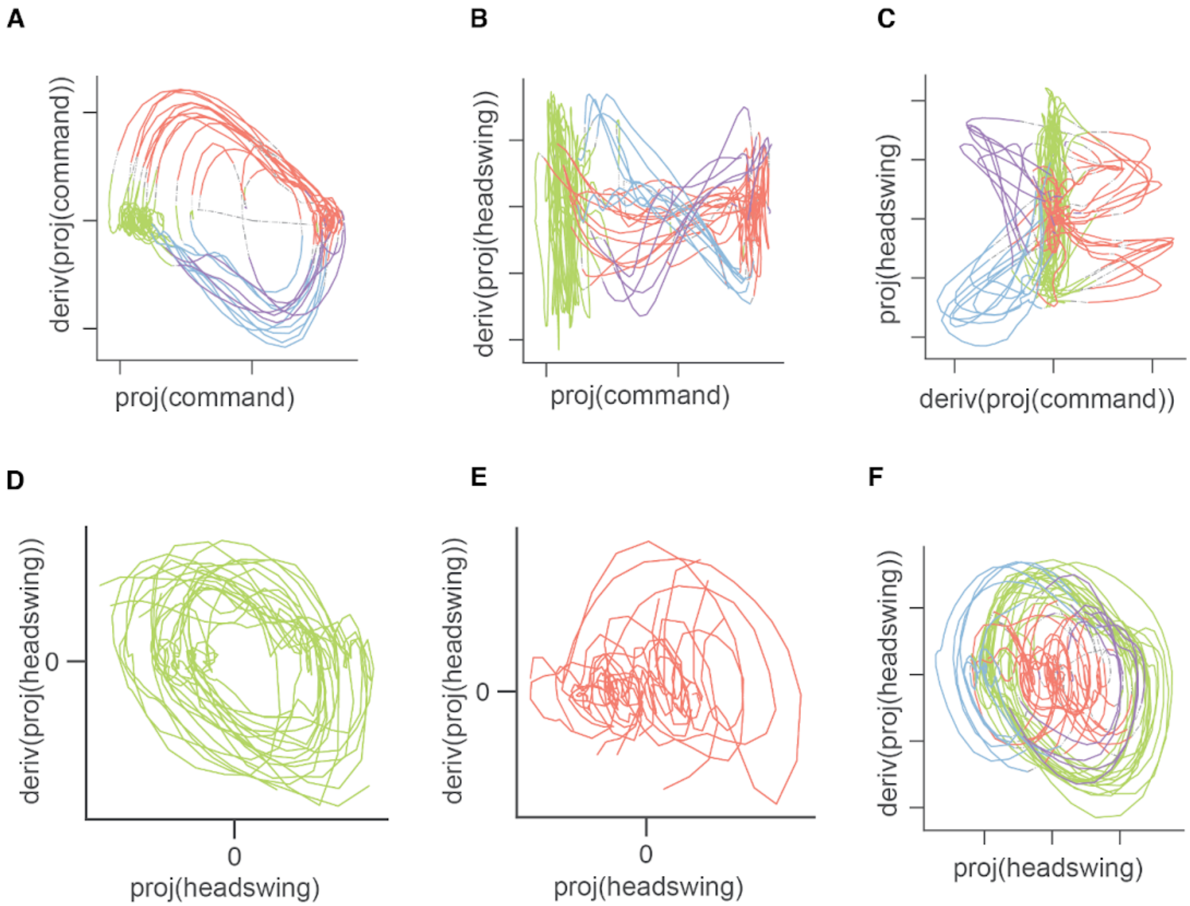


**Fig. S2. Example state space plots of projection dynamics. (A-F)** Various phase plots showing a single worm neural trajectory embedded in state space investigated in this study. Data is from same worm as in Fig. 1E, however different preprocessing parameters were used (see accompanying code). In each case, the trajectory color reflects the command state ascribed as in Fig. 1E. In (D) and (E), phase space plots of proj(headswing) reveal it rotates circularly, consistent with a sterertoyped phasic waveform, in both FWD and REV command states. We interpret the density at the origin in (E) to be consistent with the observation that our calculation of proj(headswing) does not correct for decreases in amplitude during early REV (Fig. 3f).

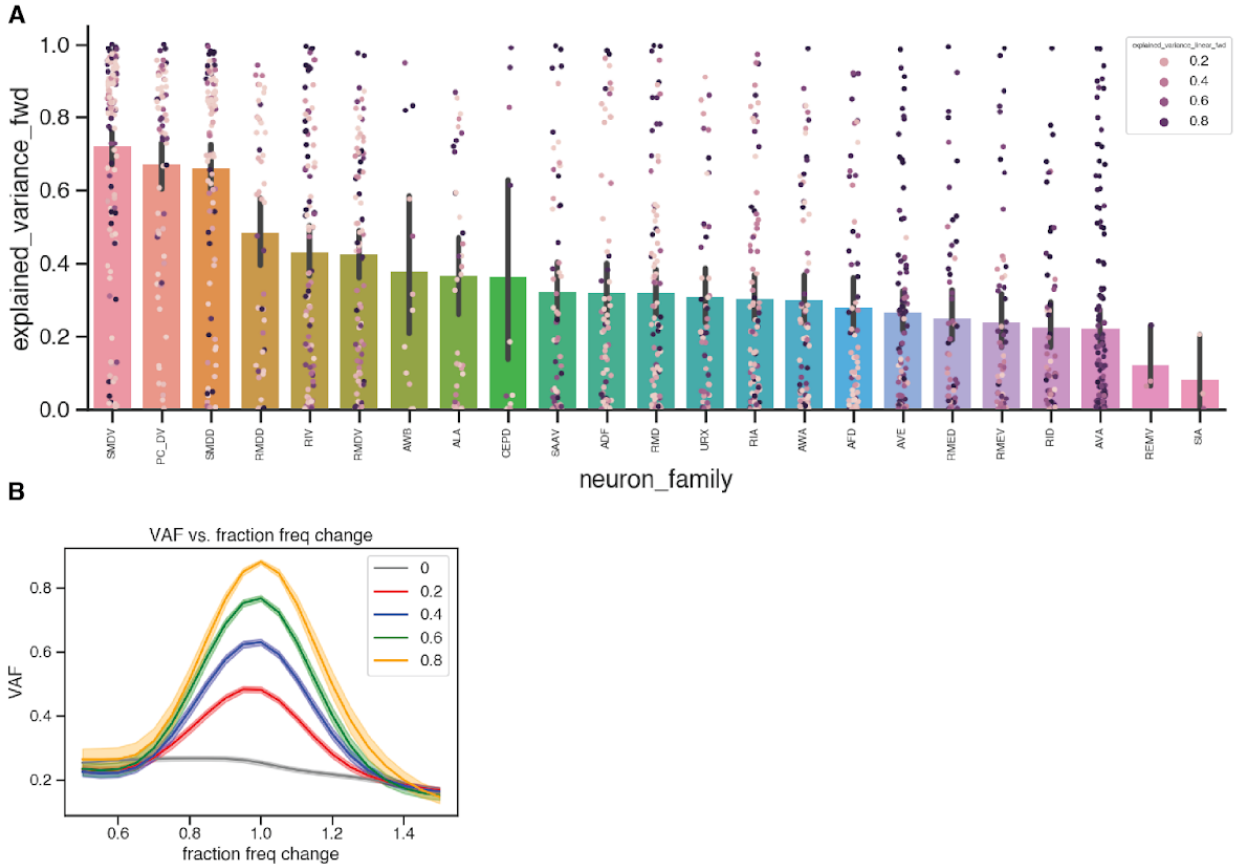


**Fig. S3. Variance explained of sinusoid fits. (A)** Percent explained variance by sinusoid fits, grouped by neuron ID. Each individual trial is plotted as a scatter point, which is colored by variance explained by a fit of a linear function. Head motor neurons exhibit both higher average VAF, as well as that VAF is not well captured by a linear fit. Error bars represent SEM. **(B)** Average variance accounted for by sinusoid fits, stratified by high-pass explained variance threshold, as a function of fit frequency, illustrating that increasing or decreasing fit frequency explained less variance of the neuron traces.

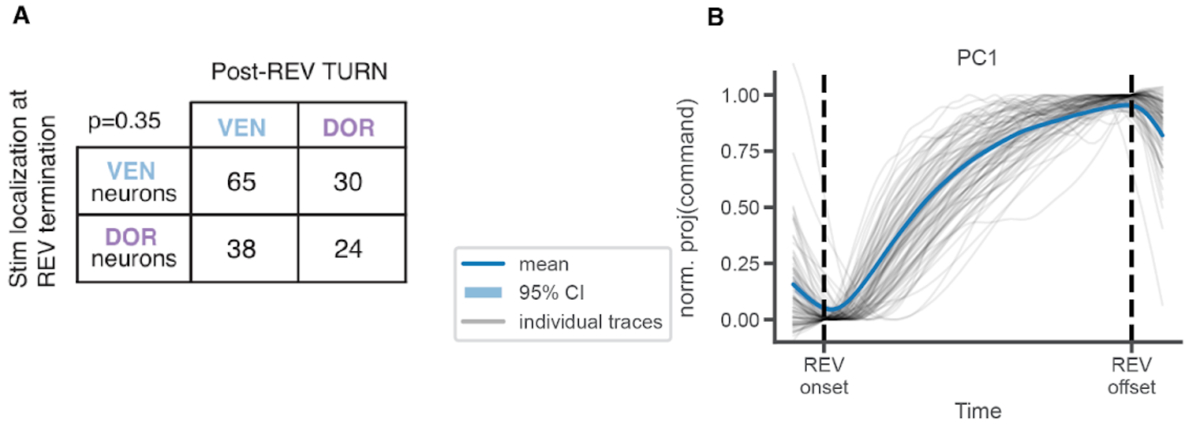


**Fig. S4. Effect of DV neuron stimulation on TURN and command state ramping. (A)** Aggregate TURN data from individual neuron stimulations in Fig 4, categorized as VEN or DOR based on whether activity of that neuron correlates with ventral or dorsal head curvature or turns. **(B)** Proj(command) normalized in amplitude and time, from REV onset to REV termination. Mean proj(command) is plotted as a thick line, and shaded regions represent 95% CI.

**Table S1. List of strains of *C. elegans* used for this study.**

| Strain name (internal) | Strain name (external) | Experiment | Genotype | Construct | Ref |
|---|---|---|---|---|---|
| FC111 | OH16230 | Control | otIs670 V; otIs672. | N/A | Yemini et al. 2021 |
| FC121 | FC121 | Fig. 4 | otIs670 V; otIs672. | PFC053 (flp-22p::Chrimson::sl2::mCherry (5ng/ul) + unc-122 RFP (50ng/ul)) | This study |
| FC128 | FC128 | Fig. 3 | otIs670 V; otIs672. | PFC079 (glr-3p::WOrMsChRmine::wrmScarlet (5ng/ul) + unc-122 RFP (50ng/ul)) | This study |
| FC134 | FC134 | Fig. 1b, Fig 2 | otIs670 V; otIs672. | PFC082 (odr-10p::WOrMsChRmine::wrmScarlet (2.5ng/ul ) + unc-122 RFP (50ng/ul)) | This study |

**Table S2. Table of Statistical Comparisons.**

| Figure | Subfigure | n (trials) | N (worms) | Statistical method | Test statistic | P-value | Testing level (alpha) |
|---|---|---|---|---|---|---|---|
| 1 | B | 148, 159 | 30 | shuffle | 124.14 | P = 1e-5 | 0.05 |
| 1 | M | (353, 1060) | 25 | KS test | 0.15 | 5.33e-07 | 0.05 |
| 2 | C | 189 | 35 | Paired ttest (2-sided) | -8.12 | 1.82e-09 | 0.05 |
| 2 | E, F | 213 | 39 | KS test | 0.29 | 8.84e-17 | 0.05 |
| 2 | H | 132 | 39 | Circular logistic regression | Log-Likelihood = -74.55 vs. LL-Null = -77.35, Pseudo $R^2$ = 0.036 | LLR p=0.018 | 0.05 |
| 2 | I | | | See below | | | See below |
| 2 | J | | | See below | | | See below |
| 3 | B | 78 | 16 | corr | 0.38, intercept: 14.94, r_value: 0.47 | p_value: 1.75e-05 | 0.05 |
| 3 | C | 70 | 15 | corr | Slope: 0.21, intercept: 0.16, r_value: 0.26 | p_value: 0.029 | 0.05 |
| 3 | D | 86 | 15 | corr | slope: -0.15, intercept: 3.85, r_value: -0.18, | p_value: 0.103 | 0.05 |
| 3 | F | 78 | 16 | corr | slope: 0.48, intercept: 22.89, r_value: 0.43 | p_value: 0.000102 | 0.05 |
| 3 | G | 57 | 15 | corr | slope: 0.22, intercept: 0.16, r_value: 0.45 | p_value: 0.000789 | 0.05 |
| 3 | H | 57 | 16 | corr | slope: 0.63, intercept: -0.079, r_value: 0.52 | p_value: 4.00e-05 | 0.05 |
| 3 | J | 66 | 16 | shuffle | tstat: 1.77 | p_value: 0.00323 | 0.05 |
| 3 | K | 66 | 16 | shuffle | tstat: 2.11 | 1e-05 | 0.05 |
| 4 | C | 131 | 14 | KS test | ks stat: 0.43 | ks p: 5.24e-22 | 0.05 |
| 4 | D | 45 | 7 | KS test | ks stat: 0.21 | ks p: 0.0296 | 0.05 |
| 4 | E | 74 | 10 | KS test | ks stat: 0.25 | ks p: 0.000193 | 0.05 |

| | | | | | | | |
|---|---|---|---|---|---|---|---|
| 4 | F | 112 | 14 | KS test | ks stat: 0.15, | ks p: 0.014 | 0.05 |
| 4 | G | 11 | 4 | KS test | ks stat: 0.37 | ks p: 0.0788 | 0.05 |
| 4 | I | 21 | | corr | $r^2 = 0.27$<br>m = 0.98 | P < 0.01 | 0.05 |
| 4 | K | 353 | 25 | corr | r=0.62, | p=4.81e-19 | 0.05 |

Additional comparisons include:

For all possible comparisons in 2I/2J:
shuffle vs AWA(1) at stim(0), tstat: 0.088, pval: 1e-5
shuffle vs AWA + proj(headswing) at stim(2), tstat: 0.153, pval: 1e-5
shuffle vs proj(headswing) at REV onset(4), tstat: 0.135, pval: 1e-5
shuffle vs proj(headswing) at stim + REV onset(5), tstat: 0.190, pval: 1e-5
shuffle vs proj(headswing) + proj(command) + SMD + RIA at stim(7), tstat: 0.274, pval: 1e-5
shuffle vs AVA at stim(8), tstat: 0.0086, pval: 0.0376
shuffle vs AVA at stim + rev(9), tstat: -0.00566, pval: 0.886
AWA(1) at stim(0) vs AWA + proj(headswing) at stim(2), tstat: 0.0737, pval: 2e-5
AWA(1) at stim(0) vs proj(headswing) at REV onset(4), tstat: 0.0535, pval: 2e-5
AWA(1) at stim(0) vs AVA at stim(8), tstat: -0.0722, pval: 2e5
AWA + proj(headswing) at stim(2) vs proj(headswing) at REV onset(4), tstat: -0.0202, pval: 0.00224
AWA + proj(headswing) at stim(2) vs AVA at stim(8), tstat: -0.146, pval: 2e-5
proj(headswing) at REV onset(4) vs AVA at stim(8), tstat: -0.1258, pval: 2e-5
proj(headswing) at stim + REV onset(5) vs proj(headswing) + proj(command) + SMD + RIA at stim(7), tstat: 0.0817, pval: 2e-5
proj(headswing) at stim + REV onset(5) vs AVA at stim + rev(9), tstat: -0.19, pval: 2e-5
proj(headswing) + proj(command) + SMD + RIA at stim(7) vs AVA at stim + rev(9), tstat: -0.2717, pval: 2e-5
